# Precision Medicine Screening Using Whole Genome Sequencing and Advanced Imaging To Identify Disease Risk in Adults

**DOI:** 10.1101/133538

**Authors:** Bradley A Perkins, C. Thomas Caskey, Pamila Brar, Eric Dec, David Karow, Andrew Kahn, Claire Hou, Naisha Shah, Debbie Boeldt, Erin Coughlin, Gabby Hands, Victor Lavrenko, James Yu, Andrea Procko, Julia Appis, Anders Dale, Lining Guo, Thomas J. Jönsson, Bryan M. Wittmann, Istvan Bartha, Smriti Ramakrishnan, Axel Bernal, James Brewer, Suzanne Brewerton, William H Biggs, Yaron Turpaz, Amalio Telenti, J Craig Venter

## Abstract

**BACKGROUND:** Progress in science and technology have created the capabilities and alternatives to symptom-driven medical care. Reducing premature mortality associated with age-related chronic diseases, such as cancer and cardiovascular disease, is an urgent priority we address using advanced screening detection.

**METHODS:** We enrolled active adults for early detection of risk for age-related chronic disease associated with premature mortality. Whole genome sequencing together with: global metabolomics, 3D/4D imaging using non-contrast whole body magnetic resonance imaging and echocardiography, and 2-week cardiac monitoring were employed to detect age-related chronic diseases and risk for diseases.

**RESULTS:** We detected previously unrecognized age-related chronic diseases requiring prompt (<30 days) medical attention in 17 (8%, 1:12) of 209 study participants, including 4 participants with early stage neoplasms (2%, 1:50). Likely mechanistic genomic findings correlating with clinical data were identified in 52 participants (25%. 1:4). More than three-quarters of participants (n=164, 78%, 3:4) had evidence of age-related chronic diseases or associated risk factors.

**CONCLUSIONS:** Precision medicine screening using genomics with other advanced clinical data among active adults identified unsuspected disease risks for age-related chronic diseases associated with premature mortality. This technology-driven phenotype screening approach has the potential to extend healthy life among active adults through improved early detection and prevention of age-related chronic diseases. Our success provides a scalable strategy to move medical practice and discovery toward risk detection and disease modification thus achieving healthier extension of life.

**SIGNIFICANCE STATEMENT:** Advances in science and technology have enabled scientists to analyze the human genome cost-effectively and to combine genome sequencing with noninvasive imaging technologies for alternatives to symptom-driven medical care. Using whole genome sequencing and noninvasive 3D/4D imaging technologies we screened 209 adults to detect age-related chronic diseases, such as cancer and cardiovascular disease. We found unrecognized age-related chronic diseases requiring prompt (<30 days) medical attention in 1:12 study participants, likely genomic findings correlating with clinical data in 1:4 participants, and evidence of age-related chronic diseases or associated risk factors in more than 3 of 4 participants. These results demonstrate that genome sequencing with clinical imaging data can be used for screening and early detection of diseases associated with premature mortality.

## INTRODUCTION

The near-doubling of average human life expectancy over the last 150 years is a tribute to scientific advancements in medicine and public health (1). In large part this success is the result of progress in control and prevention of infectious diseases particularly among young children. Eight-five percent of children born now in the US can expect to live to 65 years of age; and 42% will likely celebrate an 85 ^th^ birthday (1). As a result of this success the USA is facing a daunting and costly new medical challenge in the prevention (intervention) of age-related chronic diseases (1, 2).

Most age-related chronic diseases have heritability(3, 4), often are slowly progressive with symptom-free onset (5), and are associated with common risk factors (2, 6). In 2015, the estimated US cumulative mortality risk among males 50 to 74 years of age was 39%; for women, the risk was lower but still substantial at 24% (7). The causes of these deaths are similar across genders with neoplasms and cardiovascular disease accounting for about one-third each. Diabetes and related conditions, respiratory, cirrhosis and other liver diseases, and neurologic disorders account for the remaining one-third.

Few published examples demonstrate how genomics (8, 9) might be proactively incorporated into new models for medical practice, and what infrastructure will be needed to support data generation and use (10-15). We employed noninvasive advanced phenotyping approaches to detect risk of disease prior to damaging medical events. We used pedigree information, clinical grade whole genome sequencing (9); global metabolomics (12, 16); 3D/4D imaging focusing on non-contrast whole-body magnetic resonance imaging (17-19); cardiac imaging; 2-week cardiac rhythm monitoring; and standard laboratory studies. Our objective for active adults was similar to successful newborn screening using advanced mass spectroscopy technologies for early simultaneous detection of multiple life-threatening conditions (20, 21). Age-related chronic diseases associated with premature mortality are much more common among active adults than diseases targeted in newborn screening which make them good candidates for screening, but requiring a broader set of specialized tools and technologies for identification of disease risk than a single modality such as DNA sequencing. Phenotype data enhances risk detection and clarifies ambiguous sequence information in N of 1 evaluations.

## RESULTS

We enrolled 209 study participants, median age 55 yrs, range 20-98 yrs, 34.5% female, between September 10, 2015 and May 16, 2016. Twenty-one (10%) of the 209 participants were from 7 families. Selected characteristics comparing study participants to age and gender-adjusted NHANES cohort, a US population-based sample, is shown in Table 1. Routine clinical laboratory testing was obtained on 90 study participants (43%); Quantose IR (including fasting blood glucose) was obtained on 208 participants and included fasting blood glucose. Magnetic resonance imaging-based quantitative body compartment-specific fat and muscle estimation was conducted on 126 participants (60%). Some portion of the intended 2-week cardiac rhythm monitoring was completed on 140 (67%) participants; the median duration of monitoring was 5.9 days (range 0.8-14 days) (Figure 1).

**Figure 1.**
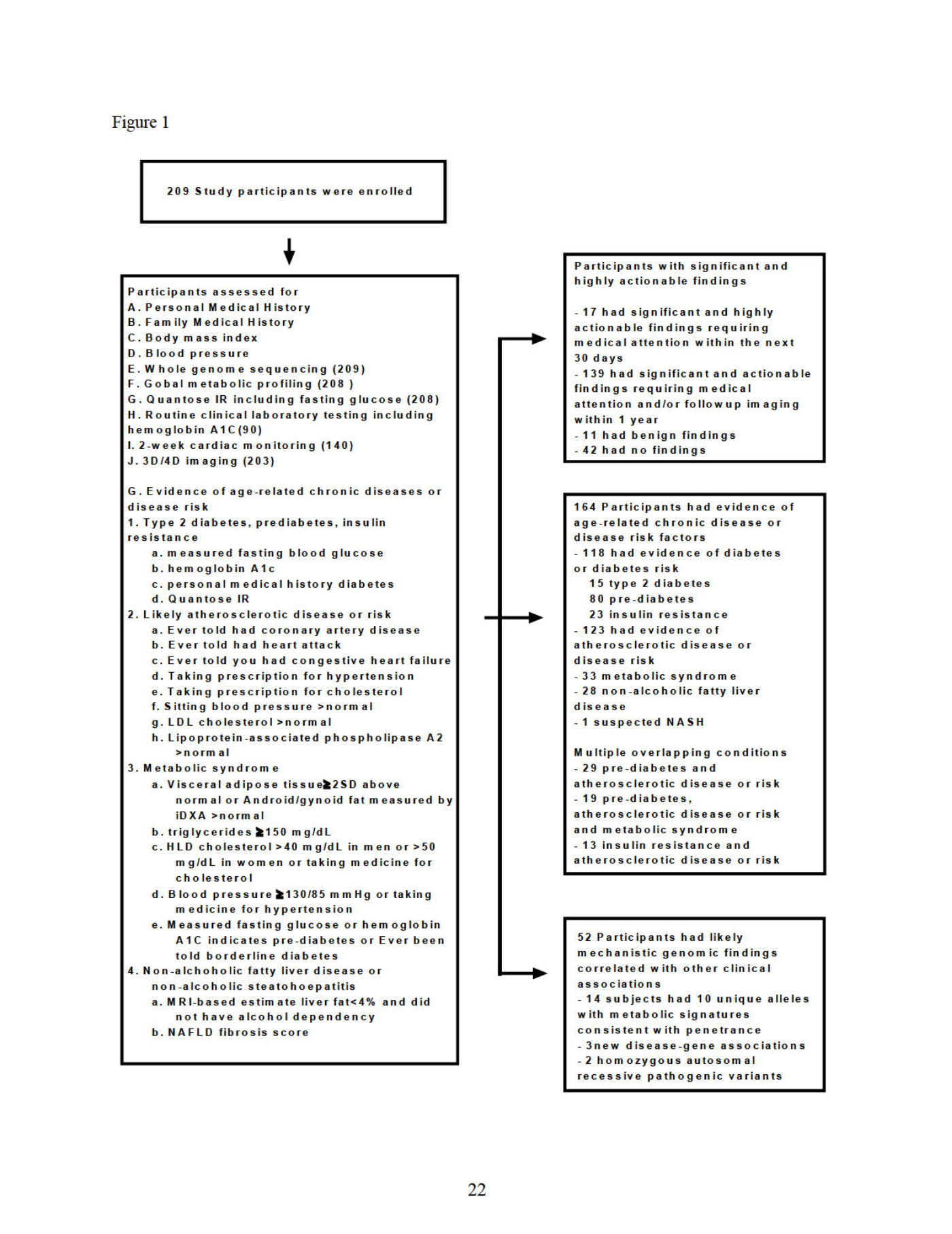
Study Design and Findings from Precision Medicine Screening.

**Table 1.** Study Participant Characteristics and Comparison to NHANES.

| Table 1. Study Participant Characteristics and Comparison to NHANES |  |  |  |  |  |
| --- | --- | --- | --- | --- | --- |
| Characteristics | Study Participant | NHANES Adult | Standardized Incidence Ratio <sup>1</sup> | 95% Confidence Interval | P-Value |
| **Age** |  |  |  |  | 4.43E-40* |
| Median | 55 | 26 |  |  |  |
| Range | 20-98 | 0-80 |  |  |  |
| **Sex** |  |  |  |  | 4.84E-04* |
| Male | 65.6% | 49.2% |  |  |  |
| Female | 34.4% | 50.8% |  |  |  |
| **Measured BMI** |  |  |  |  |  |
| Median (25%-75%) | 26 (23-29) | 24.7 (20-30) |  |  |  |
| **Measured Systolic Blood Pressure** |  |  |  |  |  |
| Median (25%-75%) | 123.5 (115-133) | 116 (106-128) |  |  |  |
| **Measured LDL** |  |  |  |  |  |
| Median (25%-75%) | 114.5 (96-135) | 103 (81-127) |  |  |  |
| **Diseases** |  |  |  |  |  |
| **Neoplasms** |  |  |  |  |  |
| Ever told you had cancer or malignancy | 15.1% | 9.5% | 1.5 | 1.02-2.16 | 3.39E-02* |
| **Cardiovascular** |  |  |  |  |  |
| Ever told you had coronary heart disease | 4.1% | 4.0% | 0.9 | 0.38-1.74 | 7.98E-01 |
| **Chronic respiratory diseases** |  |  |  |  |  |
| Ever told you had COPD? | 1.0% | 3.3% | 0.2 | 0.02-0.88 | 9.52E-02 |
| **Diabetes, urogenital, blood, and endocrine diseases** |  |  |  |  |  |
| Doctor told you have diabetes | 4.6% | 7.5% | 0.3 | 0.13-0.54 | 9.63E-04* |
| **Cirrhosis and other chronic liver diseases** |  |  |  |  |  |
| Ever told you had any liver condition | 6.1% | 4.1% | 1.1 | 0.55-1.89 | 7.75E-01 |
| **Neurological disorders** |  |  |  |  |  |
| Blood relatives have Alzheimer's disease | 13.2% | 13.3% | 1.0 | 0.63-1.44 | 1.00E+00 |
| **Risk Factors** |  |  |  |  |  |
| **Alcohol use** |  |  |  |  |  |
| Had at least 12 alcohol drinks/1 yr? | 90.0% | 70.0% | 1.2 | 0.99-1.37 | 2.76E-02* |
| **Tobacco smoking** |  |  |  |  |  |
| Smoked at least 100 cigarettes in life | 38.4% | 42.2% | 0.8 | 0.58-0.97 | 8.90E-02 |
| **High LDL cholesterol** |  |  |  |  |  |
| Now taking prescribed medicine | 78.9% | 85.4% | 1.1 | 0.74-1.48 | 6.02E-01 |
| **High blood pressure** |  |  |  |  |  |
| Ever told you had high blood pressure | 23.0% | 33.7% | 0.5 | 0.38-0.69 | 6.81E-06* |
| Taking prescription for hypertension | 73.8% | 83.6% | 0.8 | 0.54-1.14 | 2.44E-01 |
| *P ≤0.05 |  |  |  |  |  |
| <sup>1</sup> Giovanni Tripepi, 2010. "Stratification for Confounding – Part 2: Direct and Indirect Standardization". Nephron Clin Pract 2010;116:c322–c325 |  |  |  |  |  |
| NHANES, National Health and Nutrition Examination Survey, https://www.cdc.gov/nchs/nhanes/ |  |  |  |  |  |

We identified seventeen study participants (8%) with evidence of age-related chronic diseases considered significant and highly actionable requiring prompt medical attention following confirmation of screening findings: four early stage neoplasias (thymoma, renal cell carcinoma, and two high grade prostate neoplasms), one enlarged aortic root, two newly recognized atrial fibrillation cases, two medically significant arrhythmias, one 3^rd^ degree heart block, one primary biliary cholangitis, and one xanthinuria (Table 2). Some individuals had no detectable genetic risk emphasizing the value of phenotyping technology.

**Table 2.**
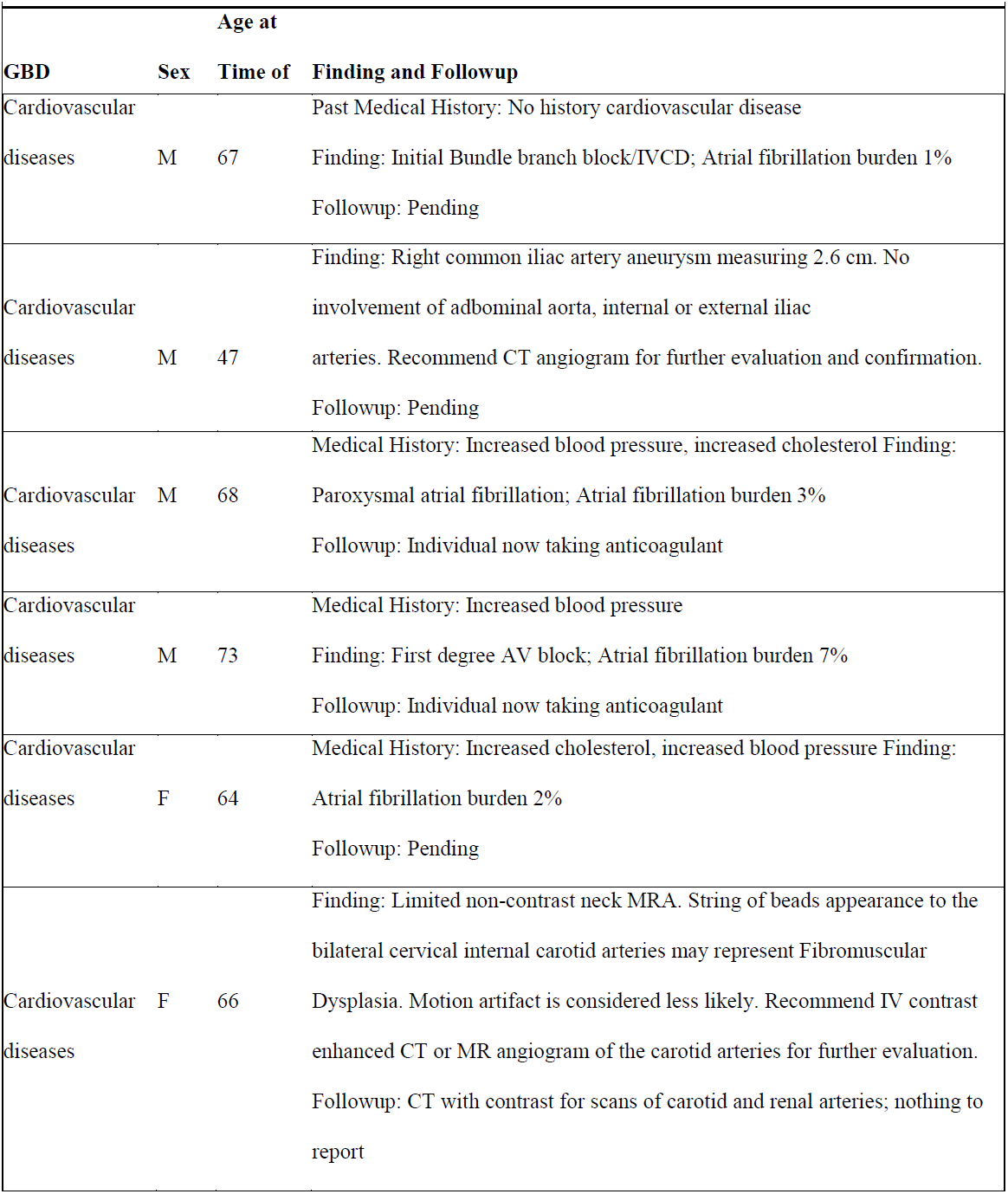
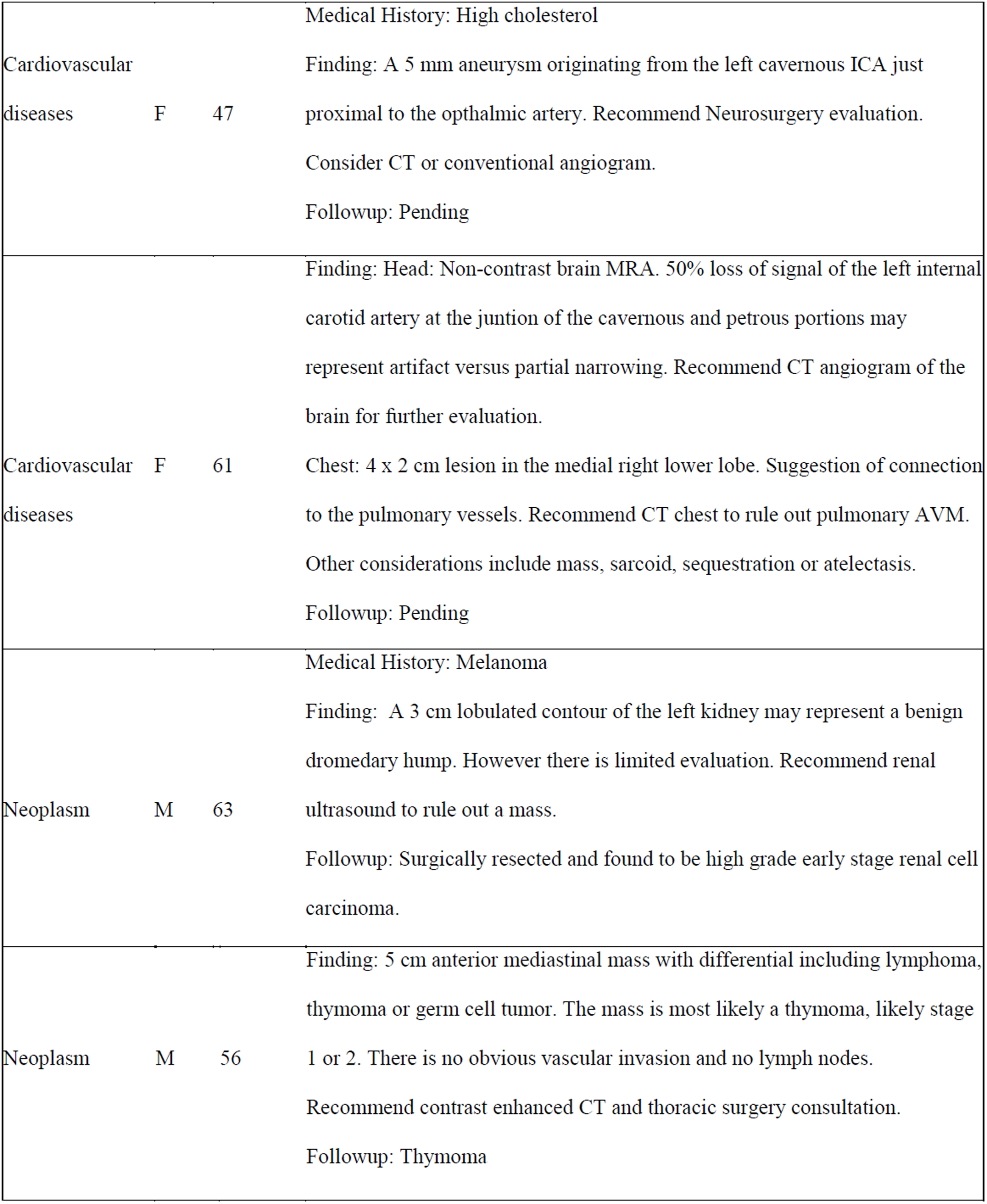
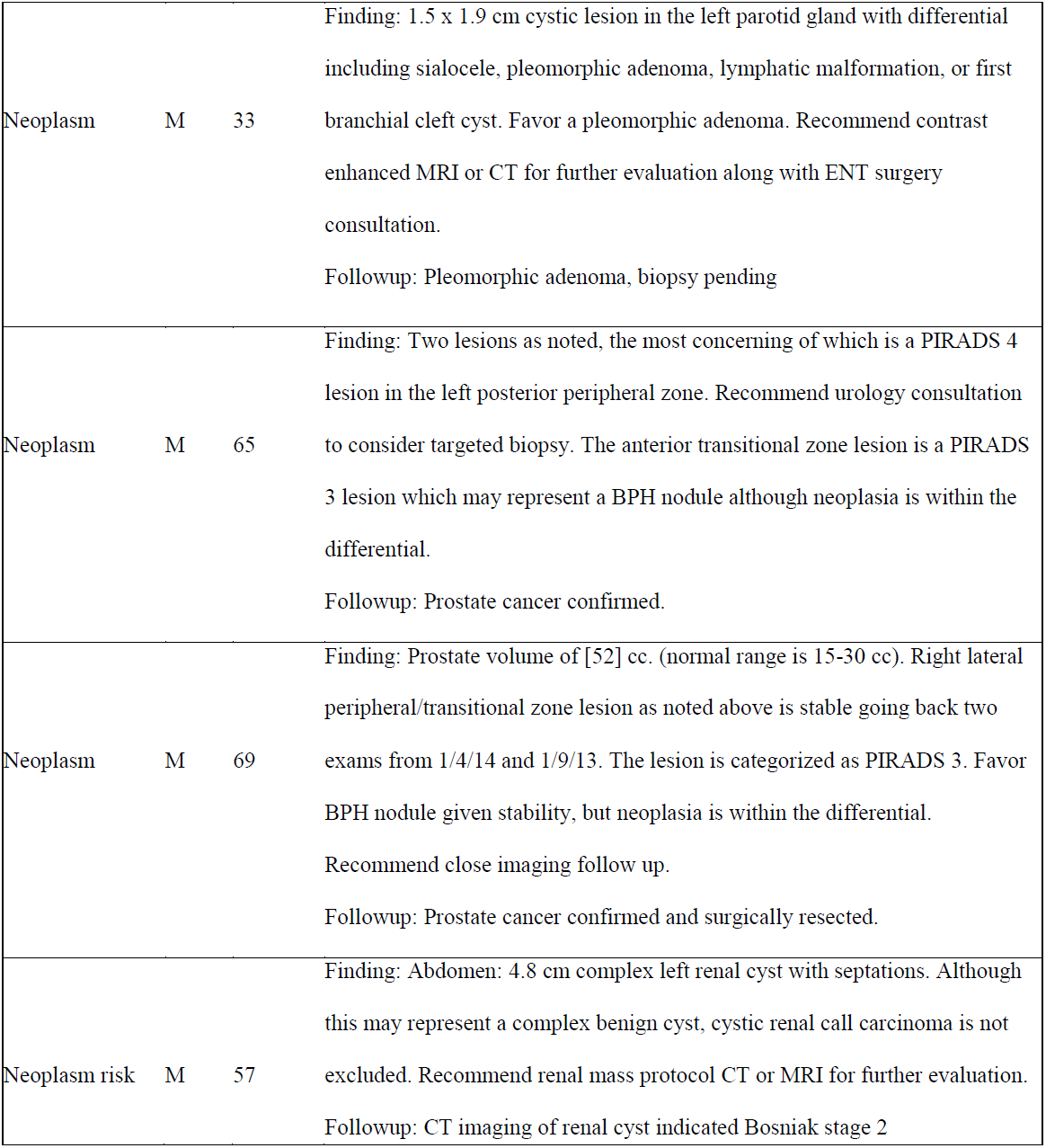
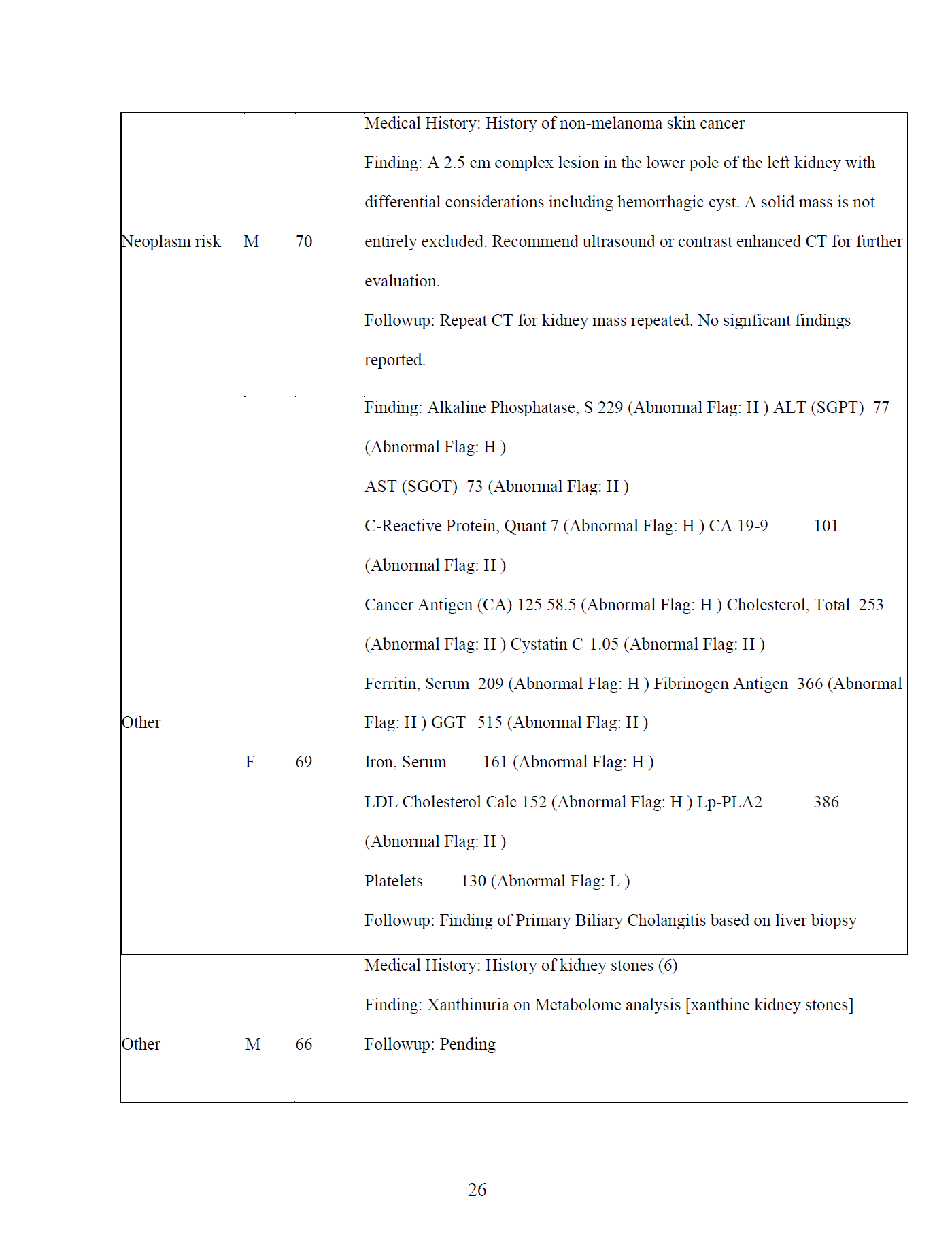
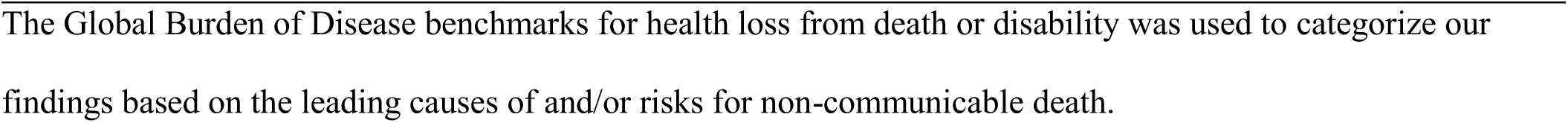
Significant and highly actionable findings.

Table 3 lists the pathogenic associations of genomic variants. 52 (25%, 1:4) participants had likely mechanistic genotype-phenotype associations (Figure 2). Of the 52 variants there were 34 unique genes, 38 unique variants, zygosity was 50 heterozygous and 2 homozygous, with 3 new variant-disease associations observed in 2 different families.

**Figure 2.**
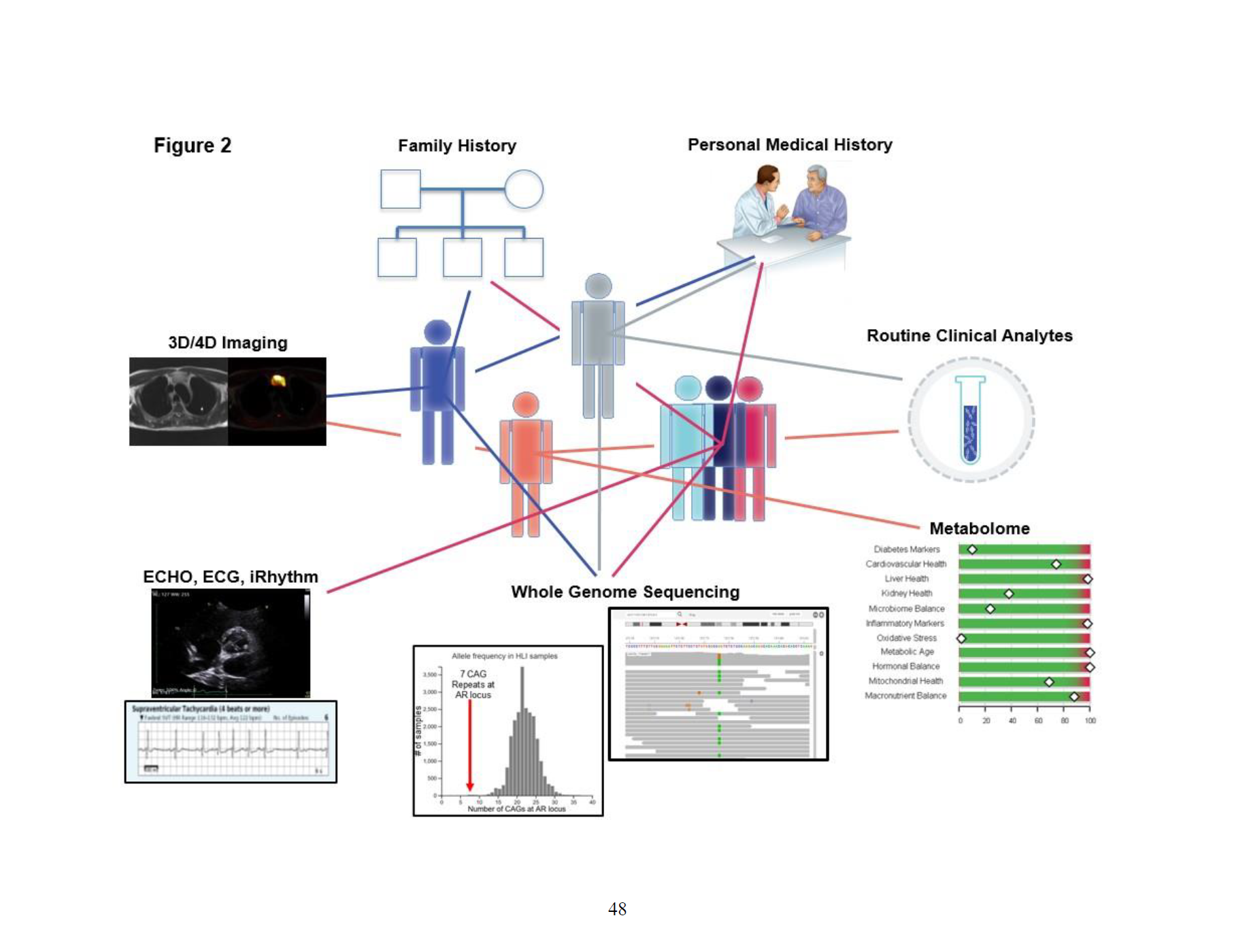
Phenotype-Genotype Data Integration. Six cases were selected to illustrate the integration of our individual technology data to achieve a precision diagnosis. Case details are found in the legend. This integration requires multiple technology skills and expert medical interpretation. Purple. Family History: 1st degree relative with two individuals with breast cancer (early onset in 40s), another first degree relative with Hodgkins lymphoma; Personal Medical History: prostate cancer diagnosed 1997, chronic lymphocytic leukemia diagnosed 2013, basal cell carcinoma and squamous cell carcinoma. Radiology: fMRI revealed focal areas of T1 hypointensity with restricted diffusion in T12, L1, L5 and S2 vertebral bodies likely hemangiomas as findings are stable; Whole Genome Sequencing: *TP53* c.844C>T (p.Arg282Trp), a likely pathogenic variant (PMID 19468865, 11370630, 8718514, 21761402, 22672556). **Gray** Family History: father with elevated cholesterol and elevated coronary calcium scoring, mother with dyslipidemia and hypertension. All grandparents had history of cardiovascular diseases; Routine Clinical Analytes: cholesterol: 247 mg/dL, triglycerides: 229 mg/dL, LDL: 157 mg/dL, VLDL: 46 mg/dL, and Lp-PLA2: 237 ng/mL; Whole Genome Sequencing: *APOB* c.9452C>T (p.Ser3151Phe), a paternally inherited rare variant. **Red** Family History: father deceased at age 83 from myocardial infarction and had a history of congestive heart failure and bundle branch block. Mother with a history of a transient ischemic attack in her 60s. Brothers and grandparents had history of high cholesterol, cardiovascular diseases or stroke. Personal Medical History: proband with dyslipidemia and noncritical coronary artery disease from calcium scoring. Cardiovascular: iRhythm showed 8 episodes of supraventricular tachycardia; Whole Genome Sequencing: a rare *DSP* c.8531G>T (p.Gly2844Val) variant (PMID 20829228) was identified in 3 siblings who also had an abnormal Personal Medical History and abnormal cardiovascular findings. **Orange** Family History: paternal grandfather with renal cell cancer, paternal grandfather’s sibling and paternal uncle with esophageal cancer. Personal Medical History: 31 yrs, BMI 33.2, a bottle of wine per day, Radiology: MRI had shown liver fat at 5%. Routine Clinical Analytes: albumin 5.0 g/dL, AST 48I U/L, GGT 111 IG/L. Metabolome: greatly reduced cysteine, cysteine sulfinic acid, 5-oxoproline and cysteinylglycine suggested that glutathione metabolism was impacted. Whole Genome Sequencing: *ALDH2* c.1510G>A (p.Glu504Lys), a pathogenic variant that had been carriers with higher acetaldehyde levels after alcohol consumption and have an increased risk of esophageal cancer (PMID 20010786).

**Table 3.** Pathogenic Genotype-Phenotype Findings.

| GBD | Gene | Disease Associated with | MOI | c.HGVS | Zygosity | Personal Medical History / Family Medical |
| --- | --- | --- | --- | --- | --- | --- |
| **Pathogenic** |  |  |  |  |  |  |
| Neoplasm | RAD50 | Hereditary cancer-predisposing syndrome | AD | c.326_329delCAGA | het | Family History: maternal family has clustering of cancer with at least 2 cases of colon cancer |
| Neoplasm | NBN | Hereditary cancer-predisposing syndrome | AD | c.657_661delACAAA | het | Family History: father has a sibling with brain cancer |
| Neoplasm | NBN | Hereditary cancer-predisposing syndrome | AD | c.127C>T | het | Family History: brother with carcinoid tumor, mother with glioblastoma multiforme, maternal aunt with breast cancer, father with amelanotic melanoma, paternal uncle with an unspecified type of cancer, paternal uncle with nasopharyngeal carcinoma |
|  |  | Aldehyde dehydrogenase deficiency, susceptibility |  |  |  | Family History: father with renal cell cancer, brother died of esophageal cancer. Primary bile acids were elevated indicative of liver dysfunction. Methionine sulfoxide and cysteine- |
| Neoplasm | ALDH2§ | Aldehyde dehydrogenase deficiency, susceptibility to esophageal cancer | AD | c.1510G>A | het | Family History: paternal grandfather died of renal cell cancer, paternal uncle died of esophageal cancer, paternal grandfather's siblings died of esophageal cancer. Liver fat: 26%, ALT is high. Extremely reduced 5-oxoproline, and cysteine suggested that glutathione metabolism was impacted. Greatly elevated cysteine-glutathione disulfide was indicative of oxidative stress. |
| Neoplasm | ALDH2§ | Aldehyde dehydrogenase deficiency, susceptibility to esophageal cancer | AD | c.1510G>A | het | Family History: paternal grandfather died of renal cell cancer, paternal uncle died of esophageal cancer, paternal grandfather's siblings died of esophageal cancer. Greatly reduced cysteine, cysteine sulfinic acid, 5-oxoproline and cysteinylglycine suggested that glutathione metabolism was impacted. Liver fat: 5%, ALT is slightly elevated. |
| Neoplasm | ATM | Hereditary cancer-predisposing syndrome, Ataxia telangiectasia | AD,AR | c.6100C>T | het | Personal History: Colon cancer; Family History: Maternal grandmother with lung cancer, father with mantle cell lymphoma and renal cell carcinoma, paternal grandmother deceased from leukemia. |
| Neoplasm | AR | Prostate cancer, susceptibility to | XLD | c.485G>A | het | Personal History: Prostate volume 52 cc. Followup imaging had predicted Gleason 4. Biopsy proven pathology bilateral Gleason 3+4. |
| Cardiovascular | PKP2 | Arrhythmogenic right ventricular dysplasia 9 | AD | c.314delC | het | Personal History: history of a mitral valve tear. ECG: Sinus bradycardia with 1st degree AV block. Rightward axis, incomplete right bundle branch block, borderline ECG. Echo: mild concentric left ventricular hypertrophy, mild enlargement of the left atrium. iRhythm: Arrhythmia: Supraventricular tachycardia, 6 episodes. Family History: father and paternal grandfather had myocardial infarction. |
| Cardiovascular | APOB | Familial hypercholesterolemia | AD | c.10580G>A | het | Routine clinical analytes: high cholesterol, LDL. Echo: mild concentric left ventricular hypertrophy. Family History: first degree relative with atherosclerosis, maternal 1st and 2nd degree relatives have cardiac problems. |
| Cirrhosis | HFE | Hemochromatosis; susceptibility to cirrhosis, diabetes and liver cancer | AR | c.845G>A | homo | Family History: sister with hereditary hemochromatosis; Personal History: possible hereditary hemochromatosis and diabetes (on metformin). iRhythm: 1 episode of ventricular tachycardia and supraventricular tachycardia; ECHO: mild concentric left ventricular hypertrophy, mild enlargement of left ventricle cavity, and a focal high signal echodensity on the aortic valve which does not have independent mobility. ECG: Right bundle branch block, left anterior fascicular block, Metabolon: impaired glucose tolerance. MRI: Liver iron level is normal (47 Hz). |
| Diabetes | SPINK1 | Pancreatitis;<br>Susceptibility to<br>fibrocalculous pancreatic<br>diabetes, Tropical<br>calcific pancreatitis | AR,AD | c.101A>G | het | Metabolic markers indicate impaired insulin sensitivity. |
| Diabetes | SPINK1 | Pancreatitis;<br>Susceptibility to<br>fibrocalculous pancreatic<br>diabetes, Tropical<br>calcific pancreatitis | AR,AD | c.101A>G | het | Metabolic markers showed impaired insulin sensitivity |
| Diabetes | SPINK1 | Pancreatitis;<br>Susceptibility to<br>fibrocalculous pancreatic<br>diabetes, Tropical<br>calcific pancreatitis | AR,AD | c.101A>G | het | Metabolic markers showed significant insulin resistance, MRI: two 6mm cystic lesions in pancreas, Routine clinical analytes: CA 19-9 is significantly high, Cancer Antigen 125 is high<br>Metabolic markers involved inflammatory are high<br>Family History: brother with diabetes |
| Diabetes | SPINK1 | Pancreatitis;<br>Susceptibility to<br>fibrocalculous pancreatic<br>diabetes, Tropical<br>calcific pancreatitis | AR,AD | c.101A>G | het | Routine clinical analytes: glucose is high<br>Metabolic markers indicated impaired glucose<br>tolerance and impaired insulin sensitivity |
| Metabolic | FMO3 | Trimethylaminuria | AR | c.458C>T | homo | Personal History: fishy odor, increased branch<br>chain amino acid metabolite markers |
| Metabolic | ACADM | Medium-chain acyl-coenzyme A<br>dehydrogenase<br>deficiency | AR | c.1084A>G | het | Medium chain acylcarnitines were greatly<br>elevated and BHBA levels low. |
| Metabolic | ACADS | Deficiency of butyryl-CoA dehydrogenase | AR | c.319C>T,c.511C>T* | het | Butyrylcarnitine and ethylmalonate were both<br>extremely elevated. |
| Metabolic | ACSF3 | Combined malonic and<br>methylmalonic aciduria | AR | c.1672C>T | het | Malonylcarnitine and 2-methylmalonylcarnitine were greatly elevated |
| Metabolic | ALDH2 | Aldehyde dehydrogenase deficiency, susceptibility to esophageal cancer | AD | c.1510G>A | het | Reduced cysteinylglycine and 5-oxoproline were suggestive of impaired glutathione metabolism. Cysteine-glutathione disulfide was greatly elevated indicative of oxidative stress. |
| Metabolic | ALDH2 | Aldehyde dehydrogenase deficiency, susceptibility to esophageal cancer | AD | c.1510G>A | het | Extremely reduced 5-oxoproline and cysteine but greatly elevated cystine suggested that glutathione metabolism was impacted. Liver fat: 5% |
| Metabolic | CTH | Cystathioninuria | AR | c.200C>T | het | Cystathionine was greatly elevated. |
| Metabolic | PAH | Phenylketonuria | AR | c.814G>T | het | Phenylalanine was high extreme and tyrosine was low. |
| **Likely Pathogenic** |  |  |  |  |  |  |
| Neoplasm | EPCAM | Lynch syndrome | AD | c.491+1G>A | het | Family History: Father with leukemia, maternal grandmother with brain tumor, maternal great aunt with liver cancer and maternal great aunt with brain cancer. |
| Neoplasm | TP53 | Osteosarcoma, Li<br>Fraumeni-like syndrome | AD | c.844C>T | het | Personal History: CLL diagnosed 2013, prostate cancer diagnosed 1997, basal cell carcinoma and squamous cell carcinoma. Family History: 1st degree relative with 2 breast primaries (early onset in 40s), another first degree relative with Hodgkins lymphoma (client believes this was acquired however). Question of Non-hodgkins lymphoma in another 1st degree relative. |
| Neoplasm | RECQL | Hereditary cancer-predisposing syndrome | AD | c.643C>T | het | Family History: mother with possibly ovarian cancer, maternal uncle with lung cancer, maternal aunt with unknown cancer, maternal uncle with possibly colorectal cancer, maternal aunt with breast cancer, maternal grandfather with unknown cancer, father with prostate and bladder cancer. |
| Cardiovascular | KCNH2 | Long QT syndrome | AD | c.2785dupG | het | Personal History: Non-specific T wave abnormality, borderline prolonged QT interval, abnormal ECG. Family History: brother with |
| Metabolic | ASS1 | Citrullinemia | AR | c.1030C>T | het | Arginine, citrulline and N-acetylcitrulline were elevated and urea was very low. |
| Metabolic | GCDH | Glutaricaciduria | AR | c.1093G>A | het | Glutaryl carnitine is extremely high |
| Metabolic | PKLR | Pyruvate kinase deficiency | AR | c.1456C>T | het | Extremely elevated glucose, elevated citrate and elevated heme possibly indicating red blood cell breakdown. |
| Metabolic | SLC7A9 | Cystinuria | AR | c.544G>A | het | Plasma cysteine extremely low. Requires urine for confirmation. |
| Other | TNFRSF13B | Common variable immunodeficiency 2 | AR,AD | c.310T>C | het | MRI: mildly enlarge periportal lymph node likely reactive (upper abdomen), spleen mildly enlarged. Routine clinical analytes: Alkaline Phosphatase, S 229 (Abnormal Flag: H ), ALT (SGPT) 77 (Abnormal Flag: H ), AST (SGOT) 73 (Abnormal Flag: H ), C-Reactive Protein, Quant 7 (Abnormal Flag: H ), CA 19-9 101 (Abnormal Flag: H), Cancer Antigen (CA) 125 58.5 (Abnormal Flag: H ), Cholesterol, Total 253 (Abnormal Flag: H ), Cystatin C 1.05 (Abnormal Flag: H ), Ferritin, Serum 209 (Abnormal Flag: H), Fibrinogen Antigen 366 (Abnormal Flag: H ), GGT 515 (Abnormal Flag: H ), Iron, Serum 161 (Abnormal Flag: H ), LDL Cholesterol Calc 152 (Abnormal Flag: H ), Lp-PLA2 386 (Abnormal Flag: H ), Platelets 130 (Abnormal Flag: L ). Finding of Primary Biliary Cholangitis based on liver biopsy. |

| Risk Factor |  |  |  |  |  |  |
| --- | --- | --- | --- | --- | --- | --- |
| Neurologic | APOE | Alzheimer's disease | AD | c.388T>C | homo | Personal History: Several mental fog recently.<br>Family History: Father with Alzheimer's disease diagnosed age 84, died of stroke age 86.<br>Paternal grandmother and grandfather with late onset Alzheimer's disease, both died in 80's. |
| VUS, VUS-suspicious |  |  |  |  |  |  |
| Neoplasm | PALB2 | Familial cancer of breast, Hereditary cancer-predisposing syndrome | AD | c.508A>G | het | Personal History: Unilateral breast cancer and unilateral ovarian cancer (Diagnosed at 32), two daughters have been through genetic test on BRCA's (negative). |
| Neoplasm | PMS1 | Lynch syndrome | AD | c.1888C>T | het | Personal History: Several colon polyps removed. Family History: possible cancers, gastric and lung, in paternal grandparent. |
| Neoplasm | CHEK2 | Hereditary cancer-predisposing syndrome | AD | c.190G>A | het | Family History: Father with prostate cancer, half brother with throat cancer, maternal grandmother had cancer twice (unknown). |
| Neoplasm | RAD50 | Hereditary cancer-predisposing syndrome | AD | c.2177G>A | het | Personal History: Breast adenocarcinoma (left), treated with lumpectomy and XRT, NED with clear nodes. Family History: Sister with leukemia at 6y. Father with metastatic lung cancer (tobacco use). Paternal aunt with breast cancer at 35y had a granddaughter with cancer at 50y, unknown type. Another paternal aunt with cancer at 63y, unknown type (tobacco use). Maternal female cousin (once removed) with skin cancer, not otherwise specified. |
|  |  | Arrhythmogenic right |  |  |  | Personal History: dyslipidemia. iRhythm: 8 episodes of supraventricular tachycardia. Family History: Father, had a pacemaker, deceased at age 83 from myocardial infarction and had a history of congestive heart failure and bundle branch block. Mother with a history of a |
| Cardiovascular | DSP¥ | Arrhythmogenic right ventricular dysplasia | AD | c.8531G>T | het | irhythm: 1 episode of supraventricular tachycardia Echo: upper limit of left ventricular wall thickness. Family History: Father, had a pacemaker, deceased at age 83 from myocardial infarction and had a history of congestive heart failure and bundle branch block. Mother with a history of a transient ischemic attack in her 60s. Paternal grandfather with likely heart attack. Paternal grandfather with likely heart attack. Maternal grandmother deceased at age 65 from a stroke. Maternal grandfather with a history of peripheral vascular disease. Maternal aunt with stroke in 50's. |
| Cardiovascular | DSP¥ | Arrhythmogenic right ventricular dysplasia | AD | c.8531G>T | het | Personal History: Hypertension diagnosed at 59y, Periodic heart flutter. ECHO: Mitral valve mildly thickened. ECG: Left atrial enlargement, borderline ECG. iRhythm: 2 episode of supraventricular tachycardia, 1 episode of ventricular tachycardia. Family History: Father, had a pacemaker, deceased at age 83 from myocardial infarction and had a history of congestive heart failure and bundle branch block. Mother with a history of a transient ischemic attack in her 60's.<br>Paternal grandfather with likely heart attack.<br>Paternal grandfather with likely heart attack.<br>Maternal grandmother deceased at age 65 from a stroke. Maternal grandfather with a history of peripheral vascular disease.<br>Maternal aunt with stroke in 50's. |
| Cardiovascular | APOB§ | Familial<br>hypercholesterolemia | AD | c.9452C>T | het | Personal History: elevated cholesterol (on Crestor) and elevated coronary calcium scoring.<br>Routine clinical analytes: Cholesterol: 237mg/dL, LDL cholesterol Calc: 154 mg/dL (range: 0-99 mg/dL)<br>Apolipoprotein (A-1): 186 mg/dL (range: 110-180 mg/dL) Apolipoprotein B: 88 mg/dL (range: 0-79 mg/dL). Family History: Mother with atrial fibrillation, hypertension and high cholesterol. |
| Cardiovascular | APOB§ | Familial | AD | c.9452C>T | het | Family History: Mother with hypertension and dyslipidemia. Father with arteriovenous malformation and high cholesterol. Paternal grandfather with cardiac valve replacement.<br>Paternal grandmother with atrial fibrillation and history of stroke, hypertension, and high cholesterol. Maternal grandmother with |
| Cardiovascular | APOB§ | Familial<br>hypercholesterolemia | AD | c.9452C>T | het | Family History: Mother with hypertension and dyslipidemia. Father with cerebralvenous malformation and high cholesterol. Paternal grandfather with cardiac valve replacement. Paternal grandmother with atrial fibrillation and history of stroke, hypertension, and high cholesterol. Maternal grandmother with vascular disease. Maternal grandfather with valvular abnormality. Routine clinical analytes: Cholesterol, total: 254mg/dL (range 100-199) LDL Cholesterol Calc: 155mg/dL (range 0-99) Triglycerides: 189 mg/dL (range 0-149) Apolipoprotein B: 147 mg/dL (range 52 -135) Apolipo. B/A-1 Ratio: 0.9 ratio units (range 0 -0.7) |
| Cardiovascular | MYBPC3 | Dilated Cardiomyopathy,<br>Hypertrophic | AD | c.1468G>A | het | ECHO: Mild concentric left ventricular hypertrophy and mild enlargement of left atrium. Family History: Father with hypertension and heart disease. Two brothers |
| Cardiovascular | MYL2 | Hypertrophic cardiomyopathy | AD | c.401A>C | het | Personal History: History of aortic valve insufficiency and cardiac enlargement; congenital bicuspid aortic valve; arrhythmia.<br>ECG: abnormal ECG. ECHO: Mild concentric left ventricular hypertrophy. Mild enlargement of left ventricle cavity. Bicuspid aortic valve.<br>Mild to moderate aortic valve regurgitation.<br>Thickened mitral valve with trace regurgitation.<br>Dilated IVC with respiratory collapse greater than 50%, consistent with mildly elevated right atrial pressure (8 mmHg). An atrial septal aneurysm is present. iRhythm: 2 episodes of supraventricular tachycardia. |
| Cardiovascular | RYR2 | Ventricular tachycardia, polymorphic | AD | c.1396C>G | het | Personal History: Cardiac palpitations history.<br>ECHO: borderline left ventricular hypertrophy.<br>ECG: Sinus bradycardia iRhythm: negative.<br>Family History: Father, brother, multiple |
| Cardiovascular | MYH7 | Cardiomyopathy | AD | c.29G>C | het | Personal History: Right bundle branch block, Echo-left ventricular hypertrophy, enlargement of left ventricle cavity, high signal echodensity on the aortic valve, suggesting focal valvular calcification. ECG: bifascicular block, right bundle branch block, left anterior fascicular block, abnormal ECG. iRhythm: 1 episode of ventricular tachycardia and supraventricular tachycardia. Family History: two maternal uncles and maternal grandmother with myocardial infarction, maternal grandfather with stroke, father with coronary artery bypass and myocardial infarction, paternal uncle with stroke, paternal grandfather with myocardial infarction. |
| Cardiovascular | LPL | Combined hyperlipidemia, familial, Lipoprotein lipase deficiency | AR,AD | c.286G>C | het | Personal History: Slightly elevated cholesterol. ECHO: left ventricular hypertrophy. ECG: possible left atrial enlargement, borderline ECG. Family History: father with hypertension, high cholesterol and heart attack at age 70, maternal cousin with heart attack in 50's and maternal grandfather with cardiovascular disease. |
| Cardiovascular | MYBPC3 | Hypertrophic cardiomyopathy; Dilated cardiomyopathy | AD | c.1000G>A | het | Personal History: Hypertension. ECHO: borderline left ventricular hypertrophy, mild regurgitation in mitral valve. Family History: mother with hypertension and arrhythmia and father with valvular heart condition. |
| Diabetes | PRSS1§ | Hereditary Pancreatitis | AD | c.107C>G | het | Personal History: type 2 diabetes mellitus, metabolic markers indicated impaired insulin sensitivity. Routine clinical analytes: Lipase |
| Diabetes | PRSS1§ | Hereditary Pancreatitis | AD | c.107C>G | het | Personal History: metabolic markers indicated impaired glucose tolerance and insulin sensitivity. Family History: mother with type 2 diabetes mellitus, maternal grandmother with a history of diabetes and maternal aunt with pancreatic cancer. |
| * NGS assay could not determine whether c.319C>T and c.511C>T occur in cis or in trans |  |  |  |  |  |  |
| § Family members with novel variant identified; ¥ Family members with novel variant identified. |  |  |  |  |  |  |

We identified 164 (78%, >3:4) participants with evidence of age-related chronic disease or risk factors. One-hundred-and-eighteen study participants (56%) had evidence of diabetes or risk for diabetes: 15 (7%) had type 2 diabetes; 80 (38%) had pre-diabetes (38%), and 23 (11%) had insulin resistance (based on Quantose *IR*). Only 19 (16%) reported a history of type 2 diabetes or pre-diabetes (Table 2). One-hundred-and-twenty-four participants (59%) had evidence of atherosclerotic disease or risk. Thirty-three (16%) had evidence of metabolic syndrome. Twenty-eight participants (13%) met a screening definition for non-alcoholic fatty liver disease (NAFLD), and one had suspected non-alcoholic steatohepatitis (NASH). Many participants had multiple over-lapping conditions including: 29 with pre-diabetes and atherosclerotic disease or risk; 19 with pre-diabetes, atherosclerotic disease or risk, and metabolic syndrome and; 13 with insulin resistance and atherosclerotic disease or risk (Figure 1).

We identified 10 unique alleles in 14 subjects with metabolic signatures consistent with penetrance. Metabolic pathways impacted by the allelic differences included fatty acid beta oxidation, fatty acid synthesis, urea cycle, and signatures associated with oxidative stress. Strong metabolic signatures were observed for two polymorphisms matching the genes’ function. Two heterozygous ACADS variants, c.1510G>A and c.1030C>T, coding for the short-chain acyl-Coenzyme A dehydrogenase (SCAD) were detected in one case. In another case, the heterozygous ACADM variant c.1456C>T coding for medium-chain acyl-Coenzyme A dehydrogenase (MCAD) was detected and interestingly both enzymes participate in fatty acid beta-oxidation by reducing different fatty acid chain length (22). SCAD specifically acts on the short chain fatty acid butyryl-CoA and MCAD reduces acyl-CoA chains containing 6-12 carbons. In the absence of SCAD activity, byproducts of butyryl-CoA including butyrycarnitine and ethylmalonate accumulate (23). Greatly elevated levels of butyrylcarnitine and ethylmalonate (Z-scores above 97.5^th^ percentile) were observed in the plasma suggestive of combined metabolic penetrance of these variants. Moreover, greatly elevated medium chain acyl-carnitines, hexanoylcarnitine, octanoylcarnitine and decanoylcarnitine (Z-scores above 97.5 the percentile) were detected suggestive of reduced MCAD activity. Large genome-wide association studies combined with metabolic profiling have previously identified associations between ACADS and MCAD and their respective metabolic substrates lending support to the metabolic penetrance observed on an individual basis in this study (24-26). We previously reported on additional metabolomic/genetic variants which are heterozygotes for known recessively inherited disorders(12, 16). These studies established that “carrier” disease state does not reflect carrier for individual metabolic variation. The number of adult cases of metabolic penetrance will continue to expanded using this approach.

Metabolomics analysis also detected xanthinuria in an individual with early onset (20’s) recurrent renal stones (6 episodes) as well as the drug effect of xanthine oxidase inhibitors in 3 other individuals. Although hypoxanthine and especially xanthine levels were elevated in both cases, normal urate and elevated orotate and orotidine levels, due to perturbed pyrimidine synthesis (27), were only observed in individuals taking xanthine oxidase inhibitors (allopurinols) for their gout conditions.

## DISCUSSION

We used a precision medicine screening approach integrating whole genome sequencing and phenotype assessments for disease risk detection among active adults focusing on age-related chronic diseases associated with premature mortality. We found a substantial burden of largely unrecognized disease risk among study participants using three different analytic perspectives including those: 1) with significant and highly actionable conditions requiring prompt medical attention for previously unrecognized potentially life-threatening age-related chronic diseases (8%); 2) with likely mechanistic genomic findings correlated with other clinical associations (25%) and; 3) with evidence of age-related chronic disease or risk factors (78%). In design of our study we hypothesized that by proactively combining clinical grade deep whole genome sequencing(9), and advanced clinical testing including global metabolomics, as well as 3D/4D imaging, emphasizing use of radiation-free low-to-no risk technologies like non-contrast MRI and echocardiography, 2-week cardiac monitoring, as well as routine laboratory testing, we could identify more precise disease risks allowing for earlier intervention and better health outcomes. False positives and other negative aspects of screening were mitigated by the high prevalence and life-threatening nature of targeted conditions, use of low-to-no-risk technologies, and convergent approaches for interpretation of results. Our data supports clinical utility of the approach presently and sets a challenge for the future improvements, including decreasing costs, through use of supporting technologies and infrastructure.

There is warranted concern about testing performance whenever screening is undertaken in medical practice. False positives may expose people to unnecessary risks, anxiety, costs, and inconvenience (28). The traditional medical approach to minimizing false positives is to rely on occurrence of symptoms to increase pre-test probabilities, though this is poorly understood by most physicians (29). Whole genome sequence data is a particular concern with the high number of VUS variants due to lack of N of 1 phenotype correlations. Traditional medical evaluations are clearly an inadequate approach for early recognition of age-related chronic diseases, many of which are preventable, and the fact that the later manifestations of these diseases now represent most of the current total US Medicare expenditure (2, 30). For nationally-sanctioned proactive single-disease adult screening programs, there are robust long-term evaluations examining testing performance in the context of clinical harms and benefits, and costs – at the population level, even though it is now increasingly well recognized that individual risk varies widely for these conditions (31). Both of these time-honored approaches have advanced health but are insufficient to cope with introduction of genomics and other new science and technologies (e.g., imaging and metabolomics) to medicine, particularly when combined with the dramatic demographic and epidemiologic changes underway in the US and globally. A major promise of genomics and precision medicine is to more tightly link curative (to identify pathology) and preventive (to identify risk) medical disciplines by creating new health and health care platforms to personalize disease risk and longitudinal care. Our data suggest a route to creating such an approach, initially focusing on prevention of premature deaths among active adults associated with age-related chronic diseases, then expanding to other causes of disability (e.g., disability-adjust life year, or DALY) and additional life stages.

Genomics as currently applied has been disappointing in its ability to unravel the “missing heritability” of most age-related chronic diseases, and other common diseases (32, 33). This shortcoming is slowly improving as a result of public and private efforts to expand sequencing but still leaves a plethora of VUS to assess. All are dependent on heterogenous contributions to the public databases, not N of 1 studies. First, we expect and are increasingly finding and seeing supporting evidence for the increasing recognition of rare variants with large effect sizes (3, 9, 34). Combining this with advancements in monogenic and polygenic methodologies to assess causation including Mendelian randomization methods (35), extension of genome-wide association study to create hazard models (36), and continued exploration of pleiotropy (37), will increase clinical utility. Second, increasingly detailed mapping of molecular pathways and mechanisms associated with diseases and risk factors will provide a much needed improved capability to link genotype and phenotype data (12, 16, 38). In our study, we were able to demonstrate the use of global metabolomics in mapping to genomic defects. This integration will strengthen with additional automation of analysis. Thirdly, we are working to quantitatively integrate genomics with other clinical data, particularly advanced imaging data, to create point-of-care clinical decision support (37, 39, 40). The version of HLI Search ^TM^ we are using can query more than 40,000 genomes (individuals and families) and explore genotype-phenotype associations with millisecond response times. We are expanding this capability using the full range of genome- and phenotype-derived data signals to rapidly identify, classify, and prioritize individual opportunities for tertiary (disease treatment), secondary (risk factor control), and primary prevention using human- and machine-driven feature extraction. Such will enable the practicing physician to incorporate precisions medicine toward disease prevention. The potential value of evolving medical practice from disease diagnosis to risk detection is supported by our study. A larger cohort and longer follow-up will strengthen these initial 209 case findings. The relatively high burden of risk identified may be partially due to self-selection of clients with health concerns. This is supported by comparison of our study participants to NHANES data (Table 2), and has been suggested by earlier authors in similar studies (11). We recommended follow-up imaging studies for slightly more than one-third of our study participants. Some of this is the nature of screening, which drives need for more definitive imaging studies better suited to specific abnormalities. Other instances of referral were intended to identify change over a specified time period which might be suggestive of cancer such as finding a cystic pancreatic lesion (41) or instability of a vascular lesion such an intracranial aneurysm (42). In some instances, data is lacking to confidently predict the natural course of these findings, and as a result may cause unnecessary anxiety and unneeded surgery (41, 42). However, the life-threatening consequences and relatively high prevalence of diseases associated with these lesions suggests that early recognition is likely to be beneficial for most individuals. Some of the technology used in this study has the potential to measure progression or resolution of the risk thus enhancing disease intervention. Expansion of some or all of our approach reported here to broader populations will require attention to cost, utility, physician and consumer acceptance. We continue to explore additional innovative noninvasive phenotyping technologies in the search for N of 1 precision medicine.

## MATERIALS AND METHODS

We enrolled active adults ≥18 years old (without acute illness, activity-limiting unexplained illness or symptoms, or known active cancer) able to come for 6-8 hours of on-site data collection, were able to undergo magnetic resonance imaging without sedation, in the case of women were not pregnant or attempting to become pregnant, and were interested in undergoing a novel precision medicine screening approach for disease risk detection including genomics and other testing, as part of an institutional review board-approved clinical research protocol. Study results were returned to study participants who were encouraged to involve their primary care physicians.

Participants underwent a verbal review of the institutional review board-approved consent (Western Institutional Review Board) and were given time to ask and receive answers to questions during a one-half to one-hour sessions conducted by health professionals. Study participants underwent standardized activities related to data collection and return of results in pre-visit, visit, and post-visit phases during a 1-year study period.

Selected data were collected regarding past medical and family history, risk factors, and medical symptoms prior to or during study participant visit (43). Participants were instructed to stop taking supplements for 72 hours, and to fast after dinner the night before their morning appointment. On the day of visit, blood was obtained for whole genome sequencing (Human Longevity, Inc.)(9), global metabolomics (12) and QUANTOSE^™^ *IR* (44) (Metabolon), and routine clinical laboratory tests (LabCorp Inc.^™^).

Two-week cardiac rhythm monitoring (Zio XT Patch^™^, iRhythm Technologies, Inc. ^™^) kits were provided with instructions for use, or monitoring was initiated during visit. Height, weight, and sitting blood pressure (45) were obtained.

Genomic variants were annotated using integrated public and proprietary annotation sources in the HLI Knowledgebase ^™^ including ClinVar(46), and HGMD ^™^ (Qiagen). Monogenic rare variants were classified as pathogenic (P), likely pathogenic (LP), or variant of uncertain significance (VUS). The HLI Knowledgebase ^™^ integrates allele frequencies for variants derived from HLI’s database of >12,000 sequences and provides a platform for query of these variants with annotation data.

To identify potentially medically significant rare monogenic variants we used an internal version (release 0.27) of HLI Search ^™^ in a two-step process: the first step focused on allele frequency <1% in the HLI cohort with annotation using ClinVar and HGMD as well as predicted loss of function variants; the second step focused on participant-specific phenotype-driven queries using an allele frequency of <1% based on family and individual medical history as well as abnormal clinical testing results.

Global metabolic profiling was performed using ultrahigh performance liquid-phase chromatography separation coupled with tandem mass spectrometry to assess the metabolic penetrance of the variants in these subject (44). Z-scores were calculated for all metabolites in each subject against a reference cohort consisting of 42 fasted subjects of normal health, and metabolites with Z-scores below the 2.5th or above the 97.5th percentiles of the reference cohort were considered to be potentially indicative of metabolic abnormalities that warranted further investigation. Integration of metabolomic and gene sequence data was achieved by a proprietary pathway analysis program developed by Metabolon and HLI.

Study participants underwent whole body magnetic resonance imaging (GE Discovery MR750w 3.0T ^™^) in research mode (courtesy GE ^™^) using protocols and post-processing for volumetric brain imaging (Neuroquant ^™^, CorTechs Laboratories ^™^), cancer detection (using restriction spectrum imaging), neurovascular and cardiovascular visualization, liver-specific fat and iron estimation, and quantitative body compartment-specific fat and muscle estimation (AMRA ^™^)(19); other post-processing was done by MMIS ^™^ (co-author, Anders Dale). GE Lunar iDXA with Pro Package was used for skeletal and metabolic health assessment. Magnetic resonance imaging and iDXA images were interpreted by co-author, DK. GE Vivid E95 was used for echocardiography and a GE Mac 2000 was used to obtain a 12-lead resting electrocardiogram. 2-week cardiac monitoring, electrocardiogram, and echocardiography were interpreted by co-author, AK.

Participants with likely mechanistic genomic findings correlating with clinical data were identified by expert review to identify convergent genomic and clinical (or phenotype) data relationships including at least two clinical (or phenotype) data elements supporting a genomic observation, including three generation family history and metabolite level correlation based on pathway mapping.

Baseline characteristics including reported past medical history for major categories of age-related chronic diseases by study participants were compared to responses from NHANES, a US population-based cohort (Table 1), adjusted for age and sex distributions. *Study participants with evidence of age-related chronic diseases considered significant and highly actionable* were defined as new genomic and/or other clinical findings which based on current medical practice indicated the need for medical attention to avoid potentially life-threatening consequences immediately or within 30 days from their visit. *Participants with evidence of age-related chronic disease or disease risk factors* were identified as including: 1) type 2 diabetes (47), pre-diabetes (47) and insulin resistance (Quantose *IR*)(43); 2) likely atherosclerotic disease or risk; 3) metabolic syndrome (48); 4) non-alcoholic fatty liver disease and non-alcoholic steatohepatitis, based on clinical guidelines or other recent literature. Measured fasting blood glucose, hemoglobin A1C, personal medical history for diabetes, or Quantose IR was used to identify participants as having diabetes, pre-diabetes or insulin resistance. The presence of any of the following were considered to be evidence of likely atherosclerotic disease or risk: “yes” in response to any of the following questions: 1) Ever told you had coronary artery disease, 2) Ever told you had a heart attack, 3) Ever told you had congestive heart failure, 4) Taking prescription for hypertension, and 5) Taking prescription for cholesterol, or if sitting blood pressure > normal, LDL cholesterol > normal, or Lipoprotein-associated phospholipase A_2_ (Lp-PLA_2_) > normal. The presence of any three of the following 5 criteria were considered to be evidence of metabolic syndrome: 1) visceral adipose tissue measured by MRI (post-processing by AMRA ^™^) ≥ 2SD above normal (19), or android/gynoid fat measured by iDXA > normal; 2) triglycerides ≥150 mg/dL; 3) HDL cholesterol <40 mg/dL in men and <50 mg/dL in women or the participant is currently taking prescribed medicine for high cholesterol; 4) blood pressure ≥130/85 mmHg or the participant is currently taking prescription for hypertension; 5) Measured fasting glucose or hemoglobin A1c indicates pre-diabetes(48) or “borderline” in response to the question - Doctor told you have diabetes. The presence of non-alcoholic fatty liver disease or non-alcoholic steatohepatitis were considered likely if: for non-alcoholic fatty liver disease MRI-based estimate liver fat was <4% and did not have alcohol dependence, and for these individuals we used a formula including other demographic and laboratory data to identify likely non-alcoholic steatohepatitis (49).

## Acknowledgements

The authors would like to acknowledge the individuals who participated in this novel precision medicine screening study without whom the findings would not be possible. Julie Ellison provided medical writing assistance, Anna Georgalis provided editorial assistance. The authors would like to acknowledge the staff past and present: Amy Reed, Ana Sanchez, Athena Hutchinson, Carina Sarabia, Cheryl Buffington, Cheryl Greenberg, Christina Bonas, Daniel Jones, Diana Cardin Escobedo, Emily Smith, Frank, Song, Genelle Olsen, Greg Olson, Heidi Millard, Helen Messier, Keisha Robinson, Laura Edwards, Nicole Boramanand, Nolan Tengonciang, Patrick Jamieson, Saints Dominguez, Samantha Punsalan, William Herrera.

## REFERENCES

1. Olshansky SJ (2016) Articulating the Case for the Longevity Dividend. Cold Spring Harb Perspect Med 6(2):a025940.

2. Bauer UE, Briss PA, Goodman RA, & Bowman BA (2014) Prevention of chronic disease in the 21st century: elimination of the leading preventable causes of premature death and disability in the USA. Lancet 384(9937):45–52.

3. Cirulli ET & Goldstein DB (2010) Uncovering the roles of rare variants in common disease through whole-genome sequencing. Nat Rev Genet 11(6):415–425.

4. Manolio TA, et al. (2009) Finding the missing heritability of complex diseases. Nature 461(7265):747–753.

5. Vogelstein B, et al. (2013) Cancer genome landscapes. Science 339(6127):1546–1558.

6. Murray CJ, et al. (2013) The state of US health, 1990-2010: burden of diseases, injuries, and risk factors. JAMA 310(6):591–608.

7. Benziger CP, Roth GA, & Moran AE (2016) The Global Burden of Disease Study and the Preventable Burden of NCD. Glob Heart 11(4):393–397.

8. Levy S, et al. (2007) The diploid genome sequence of an individual human. PLoS Biol 5(10):e254.

9. Telenti A, et al. (2016) Deep sequencing of 10,000 human genomes. Proc Natl Acad Sci U S A 113(42):11901–11906.

10. Ashley EA, et al. (2010) Clinical assessment incorporating a personal genome. Lancet 375(9725):1525–1535.

11. Gonzalez-Garay ML, McGuire AL, Pereira S, & Caskey CT (2013) Personalized genomic disease risk of volunteers. Proc Natl Acad Sci U S A 110(42):16957–16962.

12. Guo L, et al. (2015) Plasma metabolomic profiles enhance precision medicine for volunteers of normal health. Proc Natl Acad Sci U S A 112(35):E4901–4910.

13. Caskey CT, Gonzalez-Garay ML, Pereira S, & McGuire AL (2014) Adult genetic risk screening. Annu Rev Med 65:1–17.

14. Ashley EA (2016) Towards precision medicine. Nat Rev Genet 17(9):507–522.

15. Green RC, et al. (2016) Clinical Sequencing Exploratory Research Consortium: Accelerating Evidence-Based Practice of Genomic Medicine. Am J Hum Genet 99(1):246.

16. Long T, et al. (2017) Whole-genome sequencing identifies common-to-rare variants associated with human blood metabolites. Nat Genet.

17. Holland D, et al. (2009) Subregional neuroanatomical change as a biomarker for Alzheimer’s disease. Proc Natl Acad Sci U S A 106(49):20954–20959.

18. Brunsing RL, et al. (2017) Restriction spectrum imaging: An evolving imaging biomarker in prostate MRI. J Magn Reson Imaging 45(2):323–336.

19. West J, et al. (2016) Feasibility of MR-Based Body Composition Analysis in Large Scale Population Studies. PLoS One 11(9):e0163332.

20. Evans JP, Berg JS, Olshan AF, Magnuson T, & Rimer BK (2013) We screen newborns, don’t we?: realizing the promise of public health genomics. Genet Med 15(5):332–334.

21. B Therrell NNSaGRC, Austin, Texas. F Lorey, Genetic Diseases Laboratory, California Dept of Health Svcs. R Eaton, Univ of Massachusetts Medical School, Boston, Massachusetts. D Frazier, Div of Genetics and Metabolism, Univ of North Carolina at Chapel Hill. G Hoffman, Wisconsin State Laboratory of Hygiene. C Boyle, D Green, Div of Birth Defects and Developmental Disabilities, O Devine, National Center for Birth Defects and Developmental Disabilities; H Hannon, Div of Laboratory Sciences, National Center for Environmental Health, CDC (2008) Impact of Expanded Newborn Screening - United States, 2006. Morbidity and Mortality Weekly Report 57(37):1012–1015.

22. Jethva R, Bennett MJ, & Vockley J (2008) Short-chain acyl-coenzyme A dehydrogenase deficiency. Mol Genet Metab 95(4):195–200.

23. Corydon MJ, et al. (1996) Ethylmalonic aciduria is associated with an amino acid variant of short chain acyl-coenzyme A dehydrogenase. Pediatr Res 39(6):1059–1066.

24. Gieger C, et al. (2008) Genetics meets metabolomics: a genome-wide association study of metabolite profiles in human serum. PLoS Genet 4(11):e1000282.

25. Suhre K, et al. (2011) Human metabolic individuality in biomedical and pharmaceutical research. Nature 477(7362):54–60.

26. Shin SY, et al. (2014) An atlas of genetic influences on human blood metabolites. Nat Genet 46(6):543–550.

27. Beardmore TD & Kelley WN (1971) Mechanism of allopurinol-mediated inhibition of pyrimidine biosynthesis. J Lab Clin Med 78(5):696–704.

28. Weiner C (2014) Anticipate and communicate: Ethical management of incidental and secondary findings in the clinical, research, and direct-to-consumer contexts (December 2013 report of the Presidential Commission for the Study of Bioethical Issues). Am J Epidemiol 180(6):562–564.

29. Manrai AK, Bhatia G, Strymish J, Kohane IS, & Jain SH (2014) Medicine’s uncomfortable relationship with math: calculating positive predictive value. JAMA Intern Med 174(6):991–993.

30. Services CfMM (2012) Chronic Conditions among Medicare Beneficiaries, Chartbook.

31. Shieh Y, et al. (2017) Breast Cancer Screening in the Precision Medicine Era: Risk-Based Screening in a Population-Based Trial. J Natl Cancer Inst 109(5).

32. Katsanis N (2016) The continuum of causality in human genetic disorders. Genome Biol 17(1):233.

33. Manrai AK, Ioannidis JP, & Kohane IS (2016) Clinical Genomics: From Pathogenicity Claims to Quantitative Risk Estimates. JAMA 315(12):1233–1234.

34. Marouli E, et al. (2017) Rare and low-frequency coding variants alter human adult height. Nature advance online publication.

35. Ference BA, et al. (2016) Variation in PCSK9 and HMGCR and Risk of Cardiovascular Disease and Diabetes. N Engl J Med 375(22):2144–2153.

36. Ellinghaus D, et al. (2016) Analysis of five chronic inflammatory diseases identifies 27 new associations and highlights disease-specific patterns at shared loci. Nat Genet 48(5):510–518.

37. Desikan RS, et al. (2016) Personalized genetic assessment of age associated Alzheimers disease risk. bioRxiv.

38. Loscalzo J, Kohane I, & Barabasi AL (2007) Human disease classification in the postgenomic era: a complex systems approach to human pathobiology. Mol Syst Biol 3:124.

39. Hibar DP, et al. (2017) Novel genetic loci associated with hippocampal volume. Nat Commun 8:13624.

40. Zhang Z, Huang H, Shen D, & Alzheimer’s Disease Neuroimaging I (2014) Integrative analysis of multi-dimensional imaging genomics data for Alzheimer’s disease prediction. Front Aging Neurosci 6:260.

41. Konings IC, et al. (2017) Prevalence and Progression of Pancreatic Cystic Precursor Lesions Differ Between Groups at High Risk of Developing Pancreatic Cancer. Pancreas 46(1):28–34.

42. Etminan N & Rinkel GJ (2017) Unruptured intracranial aneurysms: development, rupture and preventive management. Nat Rev Neurol 13(2):126.

43. Cobb J, et al. (2013) A novel fasting blood test for insulin resistance and prediabetes. J Diabetes Sci Technol 7(1):100–110.

44. Evans AM BB, Liu Q, Mitchell MW, Robinson RJ, et al. (2014) High Resolution Mass Spectrometry Improves Data Quantity and Quality as Compared to Unit Mass Resolution Mass Spectrometry in High-Throughput Profiling Metabolomics. Metabolomics 4(132).

45. Lloyd-Jones DM, et al. (2017) Estimating Longitudinal Risks and Benefits From Cardiovascular Preventive Therapies Among Medicare Patients: The Million Hearts Longitudinal ASCVD Risk Assessment Tool: A Special Report From the American Heart Association and American College of Cardiology. J Am Coll Cardiol 69(12):1617–1636.

46. Landrum MJ, et al. (2016) ClinVar: public archive of interpretations of clinically relevant variants. Nucleic Acids Res 44(D1):D862–868.

47. Chamberlain JJ, Rhinehart AS, Shaefer CF, Jr., & Neuman A (2016) Diagnosis and Management of Diabetes: Synopsis of the 2016 American Diabetes Association Standards of Medical Care in Diabetes. Ann Intern Med 164(8):542–552.

48. Grundy SM, et al. (2005) Diagnosis and management of the metabolic syndrome: an American Heart Association/National Heart, Lung, and Blood Institute Scientific Statement. Circulation 112(17):2735–2752.

49. Angulo P, et al. (2007) The NAFLD fibrosis score: A noninvasive system that identifies liver fibrosis in patients with NAFLD. Hepatology 45(4):846–854.

